# Trophoblast organoids with physiological polarity model placental structure and function

**DOI:** 10.1101/2023.01.12.523752

**Authors:** Liheng Yang, Pengfei Liang, Huanghe Yang, Carolyn B. Coyne

## Abstract

Human trophoblast organoids (TOs) are a three-dimensional *ex vivo* culture model that can be used to study various aspects of placental development, physiology, and pathology. Previously, we showed that TOs derived from full-term human placental tissue could be used as models of trophoblast innate immune signaling and teratogenic virus infections. Here, we developed a method to culture TOs under conditions that recapitulate the cellular orientation of chorionic villi *in vivo*, with the multi-nucleated syncytiotrophoblast (STB) localized to the outer surface of organoids and the proliferative cytotrophoblasts (CTBs) located on the inner surface. We show that standard TOs containing the STB layer inside the organoid (STB^in^) develop into organoids containing the STB on the outer surface (STB^out^) when cultured in suspension with gentle agitation. STB^out^ organoids secrete higher levels of select STB-associated hormones and cytokines, including human chorionic gonadotropin (hCG) and interferon (IFN)-λ2. Using membrane capacitance measurements, we also show that the outermost surface of STB^out^ organoids contain large syncytia comprised of >50 nuclei compared to STB^in^ organoids that contain small syncytia (<10 nuclei) and mononuclear cells. The growth of TOs under conditions that mimic the cellular orientation of chorionic villi *in vivo* thus allows for the study of a variety of aspects of placental biology under physiological conditions.

## INTRODUCTION

Three-dimensional organoid culture models from tissue-derived stem cells have emerged as important *ex vivo* systems to study a variety of aspects of the physiological and pathological states of their tissues of origin. Established organoid models often preserve key features of their source organs, including tissue organization and composition, expression signatures, immune responses, and secretion profiles. Importantly, organoid cultures can be propagated long-term and can often be cryopreserved, and thus have the capacity to serve as powerful *in vitro* tools even in the absence of access to new donor tissue. Over the past several years, trophoblast organoids (TOs) derived from human placentas at different gestational stages have emerged as models by which to study trophoblast development and biology, congenital infections, and innate immune defenses^1–4^. We have shown that TOs can be derived and cultured from full-term human placental tissue and used to model trophoblast immunity and teratogenic viral infections^1^.

In tissue-derived TOs models, trophoblast stem/progenitor cells are isolated from placental chorionic villi by serial dissociation with digest solution followed by mechanical disruption (in the case of full-term tissue), then are embedded within an extracellular matrix (ECM, such as Corning Matrigel) ‘domes’. The domes containing isolated trophoblast stem/progenitor cells are then submerged in growth factor cocktail-reconstituted growth media to support stem/progenitor cell proliferation and differentiation and promote their self-organization into mature organoid units. TOs differentiate to contain all trophoblast subtypes present in the human placenta, including proliferative cytotrophoblasts (CTBs), which differentiate into the multinucleated non-proliferative syncytiotrophoblast (STB), and invasive extravillous trophoblasts (EVTs). Human chorionic villi are covered by an outermost STB layer and an inner CTB layer that fuses to replenish the outer STB during pregnancy. However, TOs cultured as three-dimensional organoids embedded in ECM develop with the opposite polarity and mature organoids contain an inward-facing STB (STB^in^) and an outward-facing CTB^1, 3^. This inverse polarity limits the utility of TOs for studies that require access to the STB layer. For example, STB^in^ TOs may not recapitulate the vertical transmission route of teratogenic infections, the transport of nutrients and antibodies across the STB, or the directionality of hormones and other factors that are critical for communication to maternal tissues and cells.

To overcome the limitation of existing TO models, we developed a suspension culture method to reverse the polarity of TOs such that the STB layer is outward facing (STB^out^). Similar approaches have been developed and applied to a variety of epithelial-derived organoid models^5–9^. We show that this culture method not only reverses the polarity of STB^in^ TOs but enhances the secretion of hormones and cytokines associated with the STB. Furthermore, we performed patch clamping of STB^in^ and STB^out^ TOs to measure the size of cells comprising the outermost layer of these organoids and found that STB^out^ organoids are covered by large syncytia (>50 nuclei), whereas STB^in^ TOs contain smaller syncytia (<10 nuclei) and are largely composed of mononuclear cells. The STB^out^ TO culture model described here thus better reflects the physiological and pathological processes of the human placenta, which can facilitate studies to define the underlying mechanisms of normal and diseased placental conditions.

## RESULTS

### Generating STB^out^ trophoblast organoids

Like epithelial-derived organoids, TOs cultured within ECM domes have an inward facing apical surface^5, 6^. However, the polarity of epithelial-derived organoids can be reversed by culturing of mature organoids under suspension culture conditions, which can occur within ∼24 hrs of initiating these cultures^5–8^. Given this, we developed a TO culturing approach that involved the culturing of organoids for 7 days in Matrigel domes to promote their differentiation and maturation, then the release of organoids from Matrigel. Once released, organoids were cultured for an additional period of 24-48 hrs in suspension with gentle agitation (schematic, **Figure 1A**). Unlike epithelial organoids in which polarity reversal can be distinguished based on brightfield microscopy alone^5, 6^, we were unable to clearly distinguish between TOs grown in suspension (STB^out^) and those cultured in Matrigel domes (STB^in^) based on brightfield microscopy alone (**Figure 1B**). To directly compare the architecture and viability of STB^in^ versus STB^out^ TOs, we performed H&E staining of cryo-sections in parallel to immunostaining these sections for a marker of CTBs (ITGA6) and the STB (CGBs). Similar to first-trimester TOs^3^, we found that STB^in^ organoids formed intracellular “mini-cavities”, with ITGA6-positive CTBs lining the outer surface of the organoids and the CGB-positive STB distributing across these intra-organoid cavities, which varied in size and quantity between organoids (**Figure 1C, top rows**). Similarly, we found that STB^out^ organoids also formed mini-cavities, but in the case of these organoids, the ITGA6-positive CTBs were largely localized to the intracellular compartment, with the CGB-positive STB on the outer surface (**Figure 1C, bottom rows**). A schematic of the general architecture of these organoids is shown in Figure 1C, left. In some cases, we did observe greater aggregation of and/or fusion between organoids in STB^out^ TOs grown in suspension (**Figure S1A**). However, this aggregation could be avoided by limiting the number of organoids seeded into each well while in suspension culture (to <100 organoids) and dissociating aggregates by manual pipetting should aggregation occurs during suspension culture.

**Figure 1:**
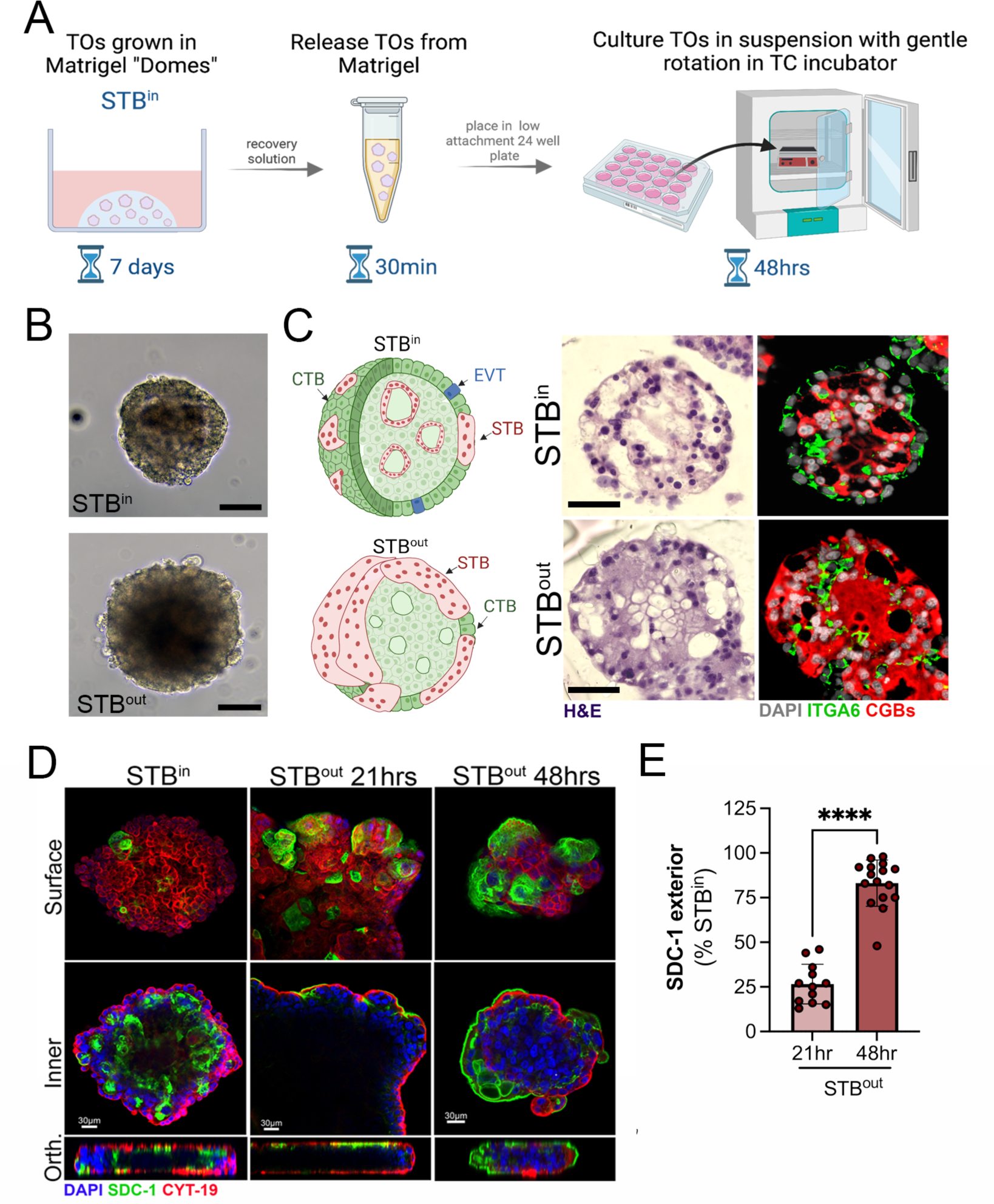
Development of STB^out^ trophoblast organoids. **(A)**, Schematic of the workflow to generate STB^out^ trophoblast organoids (TOs) from STB^in^ TOs collected from Matrigel domes. **(B),** Brightfield images of STB^in^ (top) or STB^out^ (bottom) TOs at the end of their culture period. Scale, 25μm. **(C),** Left, schematic of STB^in^ (top) or STB^out^ (bottom) TOs representing the cellular orientation of cytotrophoblasts (CTBs, in green), extravillous trophoblasts (in blue), and the syncytiotrophoblast (in red). All schematics created using Biorender. Middle, hematoxylin and eosin (H&E) histological staining of cryosections of STB^in^ and STB^out^ TOs as noted at left. Right, immunostaining for ITGA6 (green) and CGBs (red) in cryosections of matched same TOs. DAPI-stained nuclei are shown in grey. Scale, 15μm **(D),** Confocal micrographs of TOs cultured as STB^in^ (left panels) or in suspension to generate STB^out^ for 21hrs (middle) or 48hrs (right) and immunostaining for SDC-1 (green) and cytokeratin-19 (red). DAPI-stained nuclei are in blue, Top panels were captured at the outermost surface of organoids (surface) and bottom panels were captured at the innermost layers (inner). Orthogonal views (Orth) are shown at bottom. **(E),** Image analysis of the extent of surface immunostaining for SDC-1 as assessed by image analysis of SDC-1 intensity on the surfaces of TOs three-dimensional whole organoid images (shown as a percent of STB^in^ TOs) in STB^out^ TOs cultures for 21hrs (light blue) or 48hrs (dark blue). Data are shown as mean ± standard deviation with significance determined by a student’s t-test (** p<0.01). Symbols represent unique fields of organoids from individual replicates.

To determine the kinetics of the formation of STB^out^ TOs, we cultured organoids for between ∼21-48hrs and performed three-dimensional confocal microscopy using syndecan-1 (SDC-1), a cell surface proteoglycan that localizes to the apical surface of the STB, and a pan-trophoblast cytokeratin (cytokeratn-19). We found that the STB began to appear at the outer surface of TOs cultured in suspicion by ∼20hrs post-incubation but required 48hrs in culture to reach maximal exterior localization (**Figure 1D, 1E**). We did not observe any increased toxicity during this time as assessed by light microscopy, H&E staining, or lactate dehydroganse levels in media (**Figure 1B**, **1C**, and **Figure S1B**).

### Evaluation of the efficiency of generating the STB^out^ TO model at the single organoid level

To evaluate the efficiency of the STB^out^ model, and to determine whether contact between organoids was required for STB^out^ TO formation, we developed a U-bottom ultra-low attachment 96-well plate-based culture format in which individual TO unit were cultured in suspension in an individual well. To do this, we collected ∼ 10 mature TO units per donor from Matrigel domes, with lines derived from three unique placentas for a total of ∼40 organoids, in Matrigel domes. Then, individual organoids were cleared from traces of Matrigel using a needle under a tissue culture microscope and placed into a well of a U-bottom plate for 48 hr suspension culturing (**schematic, Figure 2A**, **2B, top**). Individual organoids were then fixed and immunostained for ITGA6 and SDC-1 to determine the orientation of the STB. We retrieved 37 single suspension cultured organoid units after immunostaining and found that of these organoids, 34 exhibited >50% exterior SDC-1 immunolocalization compared to matched STB^in^ organoids, in which only 1 of 37 organoids exhibited this orientation (**Figure 2C**).

**Figure 2.**
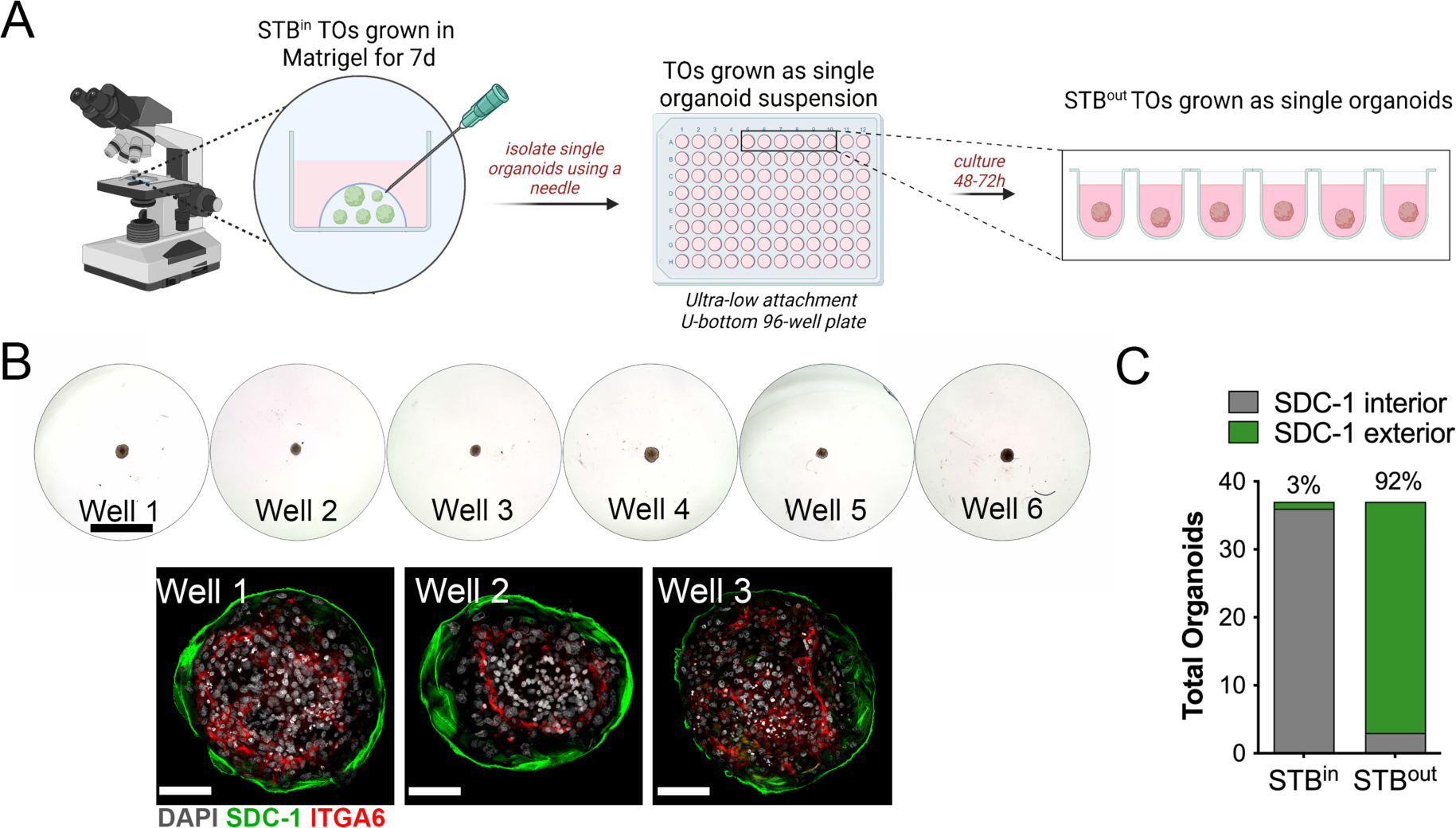
Single organoid culturing to generate STB^out^ TOs. **(A)**, Schematic depicting the workflow to generate STB^out^ TOs as single organoids in a 96-well plate format. (B), Top, brightfield images of STBout TOs cultured as single organoids. Scale bar at top, 400μm Bottom, representative images of three independent STBout TO wells immunostained for the STB marker SDC-1 (in green). DAPI-stained nuclei are shown in grey. Scale bar in bottom, 30μm. **(C),** A total of 37 organoids from three independent TO lines were cultured as single organoids and then fixed an immunostained for SDC-1 as shown in (B). Organoids with >50% SDC-1 immunolocalization on the exterior surface of the organoid (SDC-1 exterior, in green) were quantified and shown as a percentage of total organoids.

### There-dimensional imaging to define cell populations and their localization in STB^in^ and STB^out^ TOs

To further define the localization of key trophoblast cell populations between STB^in^ and STB^out^ TOs, we performed whole organoid immunostaining followed by three-dimensional confocal microscopy. SDC-1 localized to interior min-cavities in STB^in^ TOs (**Figure 3A, Supplemental Movie 1**). In contrast, SDC-1 almost exclusively localized to the outermost surfaces of STB^out^ TOs (**Figure 3A**, **Supplemental Movie 2**). Similarly, CGBs localized to the inner-most surfaces of STB^in^ TOs and were surrounded by outer layers of ITGA6-positive CTBs (**Figure 3B, Supplemental Movie 3**). In contrast, CGB localization was on the exterior of STB^out^ TOs, with ITGA6-positive CTBs comprising the interior of organoids (**Figure 3B, Supplemental Movie 4**).

**Figure 3:**
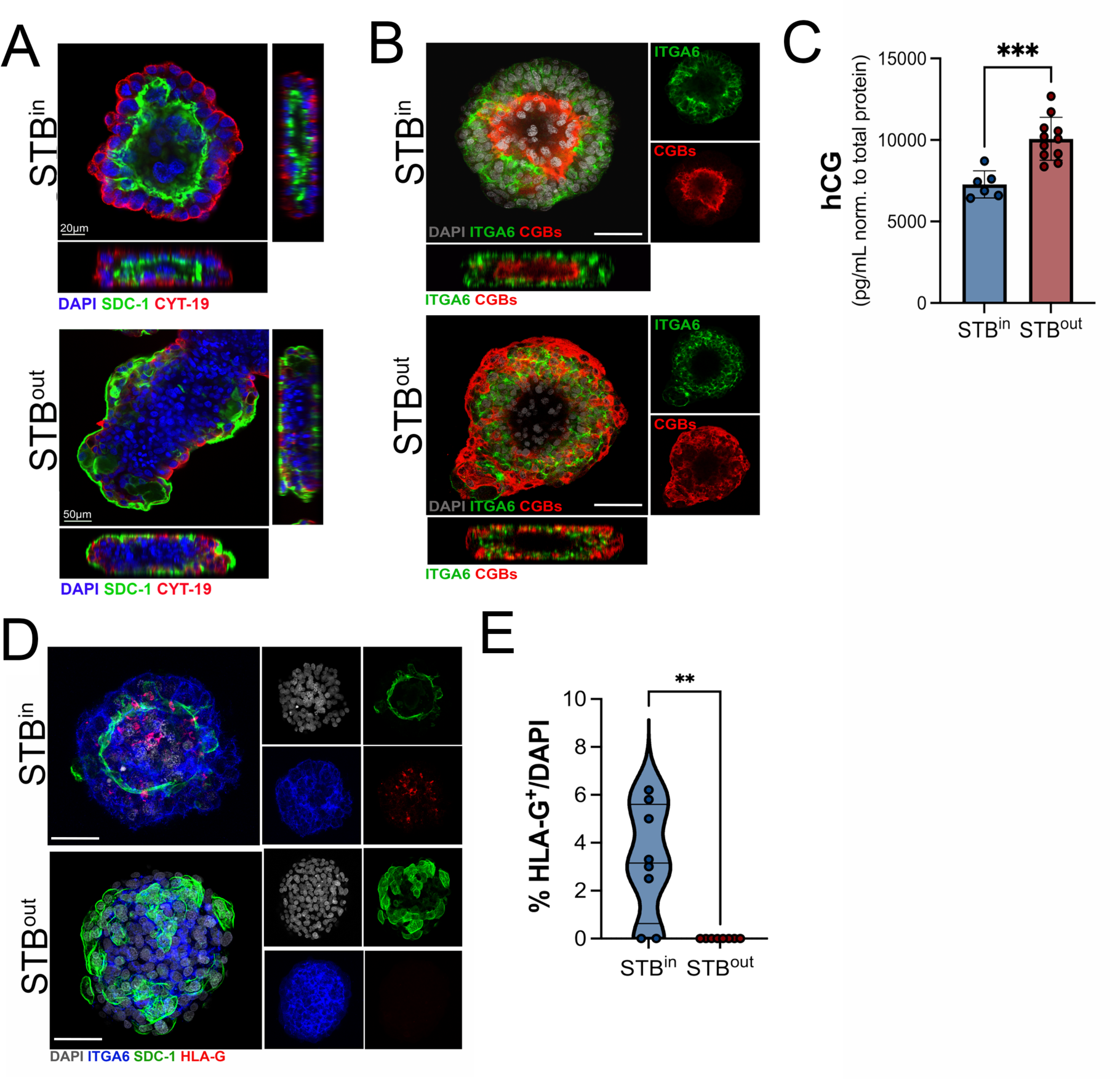
Confocal microscopy for distinct trophoblast markers in STB^in^ and STB^out^ TOs. **(A)**, Cross-sections of STB^in^ (top) of STB^out^ (bottom) TOs immunostained for SDC-1 (green), and cytokeratin-19 (red). DAPI-stained nuclei are in blue. At bottom and right are orthogonal views of three-dimensional stacked images. Movies demonstrating image reconstruction and sectioning are in Supplemental Movies 1 and 2. **(B),** Confocal micrographs of STB^in^ (top) or STB^out^ (bottom) TOs immunostained for ITGA6 (in green) or CGBs (in red). DAPI-stained nuclei are shown in grey. Right are individual channels, bottom is orthogonal views of three-dimensional stacked images. Movies demonstrating image reconstruction and sectioning are in Supplemental Movies 3 and 4. Scale bar, 30μm. **(C),** Levels of human chorionic gonadotropin (hCG) (shown as pg/mL normalized to total protein of wells from which CM was collected) in conditioned medium collected from STB^in^ or STB^out^ TOs wellsas determined by Luminex. **(D),** Confocal micrographs of STB^in^ (top) or STB^out^ (bottom) TOs immunostained for ITGA6 (in blue), SDC-1 (in green), and HLA-G (in red). DAPI-stained nuclei are shown in grey. At right are individual channels. Scale bar, 30μm **(E),** Quantification of the percentage of HLA-G^+^ positive cells in STB^in^ versus STB^out^ TOs. Individual points represent unique fields used for image analysis. A minimum of 15 organoids was used for analysis. Significance determined by a student’s t-test (** p<0.01).

The STB is a primary producer of hormones required for pregnancy, including human chorionic gonadotropin (hCG) which is comprised of two subunits, CGBs and CGA. We and others have shown that STB^in^ TOs recapitulate this secretion^1, 3^. To determine if there were differences in the secretion of hCG between STB^in^ and STB^out^ TOs, we performed Luminex assay of conditioned medium from both STB^in^ and STB^out^ TOs wells. To control for variability in organoid size and quantity, we also collected and quantified total protein used to normalize these values. We found that there were significantly higher levels of hCG in media collected from STB^out^ TOs compared to STB^in^ TOs (∼10000pg/mL versus ∼7000pg/mL, respectively) (**Figure 3C**), consistent with the enhanced localization of the STB to the outer surfaces of TOs.

TOs can self-differentiate to contain small amounts of HLA-G^+^ EVTs^1, 2^. To determine if this also occurred or was altered by STB^out^ suspension culture conditions, we performed immunostaining for HLA-G in STB^in^ and STB^out^ TOs. Consistent with our previous study, we found that STB^in^ TOs differentiated to contain <10% HLA-G^+^ EVTs (**Figure 3D, 3E**). In contrast, we were unable to detect any HLA-G^+^ cells in STB^out^ TOs, suggesting that suspension culture condition reduces spontaneous EVT differentiation. Although rates of EVT differentiation can be promoted by altering composition of culture media^1, 2^, this process requires an extended culture period of >3-4 weeks, which is beyond the time frame possible to culture STB^out^ TOs in suspension. Thus, it remains unclear if these STB^out^ organoids can also be cultured to promote EVT differentiation.

### Profiling of cytokine and chemokine secretion in STB^in^ and STB^out^ organoids

In addition to hormones, the STB also secretes cytokines required to facilitate the establishment of tolerance and/or to defend the fetus from pathogen infection, such as the release of the antiviral type III interferons (IFNs) IFN-λs^11^. We showed previously that TOs recapitulate this secretion and release a number of these cytokines, including IL-6 and IFN-λ2^1^. To determine if STB^out^ TOs maintain this cytokine secretion or induce unique cytokines and chemokines compared to STB^in^ TOs, we performed multiplex Luminex profiling of 73 cytokines and chemokines, a subset of which we previously showed were released from STB^in^ TOs^1^. We did not observe any secretion of cytokines and chemokines in STB^out^ TOs that were not also secreted from STB^in^ TOs (**Figure 4A**). However, we found that STB^out^ TOs secreted higher levels of two factors, IFN-λ2 (>7-fold increase) and IL-6 (>4fold increase), (**Figure 4B-D**). In contrast, other analytes such as GRO-α/MIF were secreted at similar levels between both STB^in^ and STB^out^ TOs (**Figure 4A, 4E**). One chemokine, CXCL1, which has been associated with decidual stromal cell responses to trophoblasts^12^ and which uses SDC-1 as a co-receptor^13^, was significantly reduced in STB^out^ TOs (>12-fold reduction) (**Figure 4B, 4F**). Collectively, these suggest that STB orientation in TOs has implications on immune secretion profiles.

**Figure 4:**
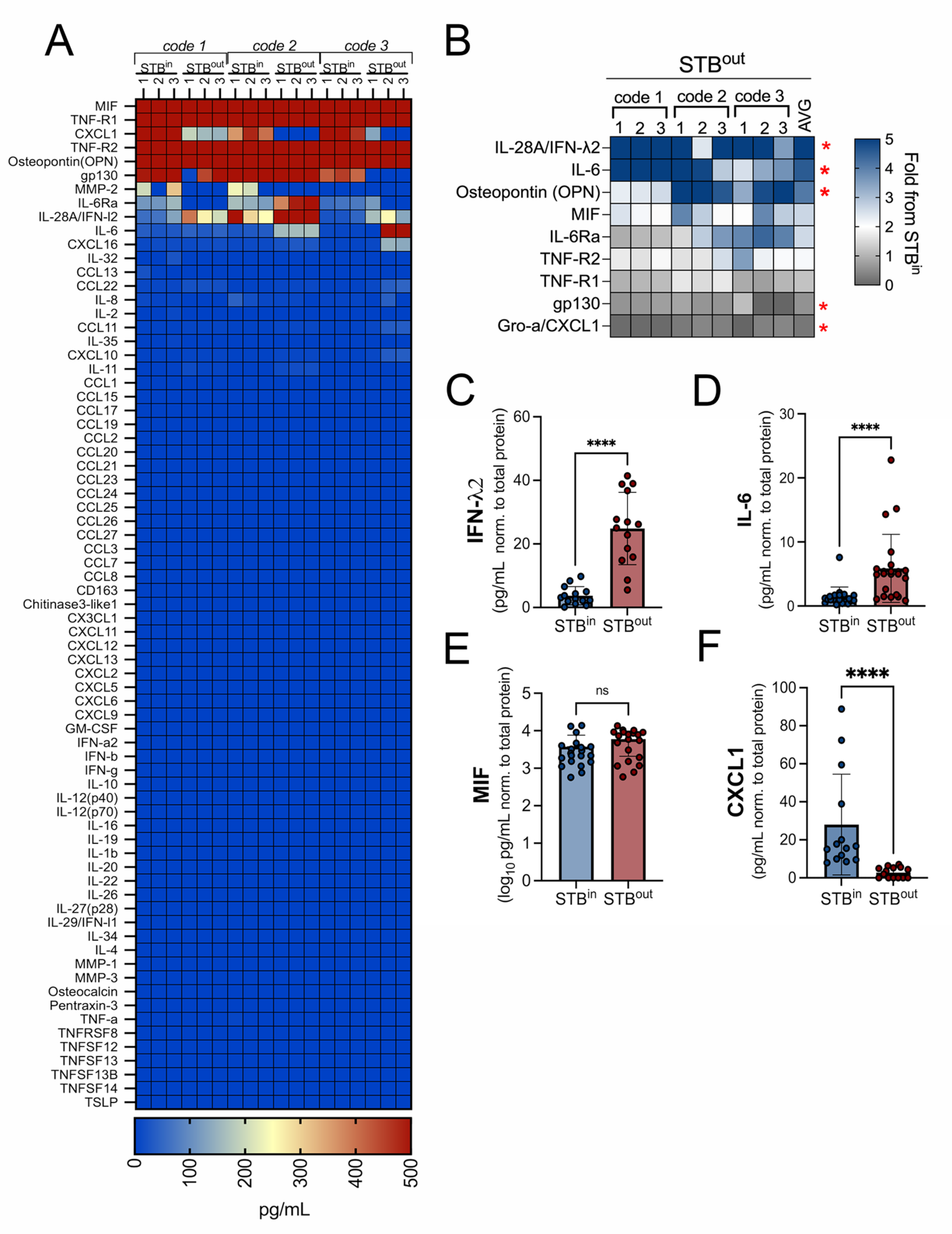
Levels of STB-associated cytokines and chemokines in STB^in^ and STB^out^ TOs. **(A)**, Heatmap depicting the levels of cytokines and chemokines in conditioned medium (CM) isolated from three independent codes of TOs derivedfrom unique placental tissue. Three replicate wells of each condition are shown. Analytes with detected values at or above 500ng/mL are shown in red and analytes with no to little detection are shown in blue, as shown in scale at bottom. **(B),** Heatmap of cytokines and chemokines released from STB^out^ TOs. Data are shown as a fold-change from STB^in^ organoids (blue is increased and grey is decreased levels). Red asterisks designate factors increased in STB^out^ TOs by >2-fold or decreased >2-fold. Data are shown from twelve independent CM preparations from at least 3 codes, with average shown at right. **(C-F),** Levels of IFN-λ2 (C), IL-6 (D), MIF (E), or CXCL1 (F) from CM collected from STB^in^ or STB^out^ organoids as determined by Luminex assays. All data are shown as pg/mL and normalized to total protein of wells from which CM was collected. All data are also shown as mean ± standard deviation with significance determined by a student’s t-test (****, p<0.0001, ns, not signfiicant). Symbols represent unique media samples collected from replicate experiments from at least three unique codes.

### Membrane capacitance measurements confirms the presence of large syncytia on the exterior surface of STB^out^ TOs

Cell fusion dramatically increases the surface area of the fused cell. As cell surface area is proportional to its membrane capacitance (Cm)^14^, patch clamp, a quantitative electrophysiological technique^15, 16^, can be used to evaluate cell size. We therefore utilized patch clamping to calculate the size of cells/syncytia comprising the exterior cellular surface of STB^in^ versus STB^out^ TOs (schematic, **Figure 5A**). When a small voltage step (10 mV) was applied to TO lines derived from three unique placentas, the capacitive current from STB^out^ TOs showed much slower decay than the capacitive current from STB^in^ TOs (**Figure 5B**). The average Cm in STB^in^ were 0.238 nF, 0.076 nF and 0.104 nF for code 1, code 2 and code 3, respectively. Whereas the average Cm in STB^out^ TOs were 2.812 nF, 1.603 nF and 1.899 nF, respectively for the three codes. (**Figure 5C**). Interestingly, the Cm of the surface trophoblasts in STB^out^ TOs exhibited a Gaussian distribution in all lines tested (**Figure 5D-F**). In stark contrast to the broader distribution of the Cm from STB^in^ TOs centered at 0.113 nF, 0.051 nF and 0.087 nF for code 1, code 2 and code 3, respectively, the Cm from the STB^out^ TOs was largely centered at 3.350 nF, 1.230 nF and 1.622 nF, about 20-30 fold larger than in STB^in^ TOs. It is worth noting that extremely large syncytia are readily observed on the surfaces of STB^out^ TOs (**Supplemental Figure 2A**). We recorded 5 independent areas of these cells and found that they have unmeasurable cell capacitance (**Supplemental Figure 2B**). This is likely due to space clamp issues for syncytia with extremely large surface areas^17^. It is noteworthy to mention that the smallest Cm measured in our study, which represents the Cm of a single cell, was approximately 30 pF. Syncytialization involves fusion of plasma membranes, and Cm is directly proportional to the cell surface area. Therefore, the extend of syncytialization in STB^in^ and STB^out^ TOs can be estimated by dividing the Cm of the fused cell by the single cell capacitance of 30 pF. By performing the necessary calculations, we found that the surface trophoblasts observed in the STB^in^ samples were predominantly composed of individual cytotrophoblasts (CTBs) and syncytia with limited fusion (less than 10 nuclei). On the other hand, the surface trophoblasts in the STB^out^ samples are primarily comprised of syncytia with a greater number of nuclei, exceeding 50 nuclei.

**Figure 5:**
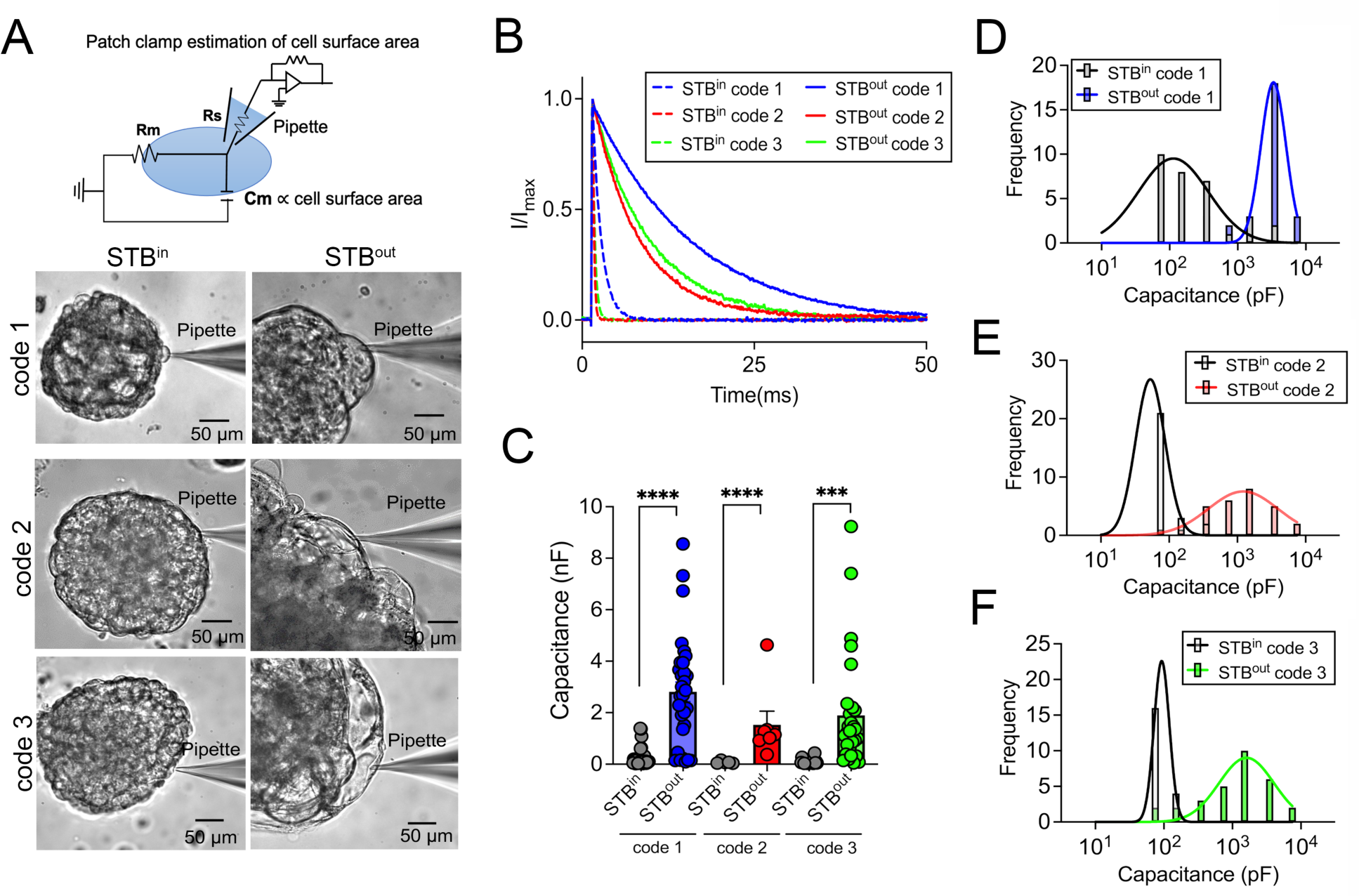
Evaluation of trophoblast fusion on the surface of STB^in^ and STB^out^ TOs using membrane capacitance measurement. **(A)**, Top, diagram of whole-cell patch clamp to measure membrane capacitance (Cm), which is proportional to cell surface area. Rs: series resistance; Rm: membrane resistance. Bottom, representative brightfield images of patch-clamped surface trophoblasts from the TOs growing under STB^in^ (left) or STB^out^ (right) conditions. **(B),** Representative membrane test traces from STB^in^ (dashed lines) and STB^out^ (solid lines) TOs to measure cell capacitance from three TO codes derived from independent placental tissues (in blue, red, and green). Current was elicited by a test voltage pulse of 10 mV from a holding potential of 0 mV (top). **(C),** Summary of membrane capacitance measured from STB^in^ (grey) and STB^out^ (blue, red, and green) TOs derived from independent placental tissues. Two-sided Student’s t-test. (n=31 for both STB^in^ and STB^out^ code 1; n=26 and 28 for STB^in^ and STB^out^ code 2, respectively; n=22 and 29 for STB^in^ and STB^out^ code 3, respectively. ***p<0.001, ****p<0.0001). **(D-F)** Distribution of cell capacitance from STB^in^ (black) and STB^out^ TOs from three TO codes derived from independent placental tissues (in blue (D), red (E), and green (F)). The bars were at the center of each bin (see Methods for details). The data were fitted with Gaussian distribution.

## DISCUSSION

In this study, we develop a method to culture trophoblast organoids under conditions that reflect their physiological cellular orientation *in vivo*. This model facilitates access to the STB layer while also maintaining key features associated with STB^in^ TOs, including their three-dimensional morphology, the presence of distinct trophoblast subpopulations, and the secretion of pregnancy related hormones and immune factors. STB^out^ TOs have several advantages over STB^in^ TOs. For example, STB^out^ TOs naturally self-reorganize with an STB outward-facing surface and do not require extensive manipulation to develop this outer layer. In addition, as STB^out^ TOs are cultured in suspension, the lack of ECM allows for applications in which this scaffold presents a barrier to diffusion, such as studies of microbial infections or antibody uptake.

For epithelial organoids grown in ECM domes with basal-out polarity, microinjection can serve as an option to directly access the enclosed apical surface^18, 19^. However, in contrast to epithelial-derived organoids which often form clear cystic structures, TOs have heterogeneous mini-cavities, which makes microinjection of these organoids difficult. Additional methods have been applied to epithelial-derived organoids, such as seeding dissociated organoid fragments onto Transwell inserts^20, 21^. However, this approach compromises the three-dimensional nature of organoids which may impact their function. The method we describe here avoids several of these challenges, as STB^out^ TOs maintain their three-dimensional structure and do not require their disruption to generate. It is unclear whether STB^in^ TOs undergo similar mechanisms of polarity reversal as do epithelial-derived organoids, which undergo relocalization of junction-associated proteins to mediate this process, or whether culturing in suspension instead promotes CTB fusion on the organoid surface. Given that the surface of STB^out^ TOs is covered by very large syncytia, it is possible that suspension culturing promotes the fusion of CTBs on the organoid surface rather than inducing a relocalization of the STB from the inner to outer organoid surface. This fusion could be promoted by factors including low levels of shear stress during suspension culturing, which has been proposed to enhance rates of CTB fusion^22^. Fluid shear is known to impact myriad aspects of epithelial cell function, including formation of microvilli^23^, suggesting that this shear likely also impacts rates of CTB fusion.

A benefit of TOs is their ability to recapitulate the hormone and cytokine secretion observed in primary trophoblasts and chorionic villous tissue explants^1^, which is not recapitulated in standard trophoblast cell lines^11^. However, given that STB^in^ TOs are embedded in Matrigel, many of these STB-associated factors would be secreted into the center of the organoid structure or perhaps into the surrounding ECM. We found that STB^out^ TOs not only recapitulate the release of these factors, but that some factors were secreted at significantly higher levels than those observed in STB^in^ TOs. The mechanistic basis for this is likely two-fold and could include the increase in syncytia size on the STB^out^ TO surface as well as the direct release of these factors into the culture media. However, we found that STB^out^ organoids exhibited substantially lower levels of HLA-G^+^ EVTs, suggesting that there is reduced EVT differentiation, which could impact EVT-specific secretion profiles. It is not clear whether methods to promote EVT differentiation previously applied to TOs derived from full-term tissue^1^ could also be applied to the STB^out^ TO system. However, given the extended time to perform this procedure (>3 weeks), it is unlikely that STB^out^ TOs would be amenable to this process.

We leveraged the power of electrophysiology to define the size of cells/syncytia covering the surface of STB^in^ and STB^out^ TOs. These studies verified the high efficiency of the STB^out^ TO system and provided quantitative measurements of the number of nuclei comprising syncytia. Based on these findings, we estimate that syncytia covering STB^out^ TOs were comprised of at least 60 nuclei as well as some syncytia that were too large to be measured by patch clamping. These studies not only confirmed the presence of syncytia on the outer surface of STB^out^ TOs but provide a strong proof of concept for the application of this approach to quantitatively measure syncytial size on the surfaces of TOs, which could be applied to a variety of biological questions.

The STB^out^ model we describe does have limitations. We have applied the protocol described above to multiple lines of TOs derived from unique full-term placental tissue. Although we would anticipate that this protocol can be adapted to TOs derived from early gestation tissue as well as from iPSC-derived organoids, it is possible that some of the steps described would require further optimization for these models. The most significant limitation of this approach is that STB^out^ organoids cannot be passaged to maintain this physiological polarity phenotype, and TOs with this orientation must be generated for each experiment. Thus, generation of STB^out^ TOs is considered a terminal culture approach, like existing EVT differentiation methods^2^. However, given that STB^out^ polarity is maintained post-culturing, these organoids can be utilized for a studies post-generation.

The model described here provides an organoid system that recapitulates the cellular orientation of the human placenta *in vivo* and provides evidence that this system can be used to model key aspects of STB structure and function. In addition, given that we have developed a system for single organoid culturing in a multi-well format, this approach could be used for mid-to-high throughput screening approaches. Collectively, our method described here can be used model key aspects of placental physiology and development.

## MATERIALS AND METHODS

### Trophoblast organoid culturing

TO lines used in this study were derived from human full-term placentas as described previously^1^. For passaging and culturing, TOs were plated into Matrigel (Corning 356231) domes, then submerged with prewarmed complete growth media as described^1^. Cultures were maintained in a 37°C humidified incubator with 5% CO2. Medium was renewed every 2-3 days. About 5-7 days after seeding TOs were collected from Matrigel domes, digested in prewarmed TrypLE Express (Gibco, 12605-028) at 37°C for 8 min, then mechanically dissociated into small fragments using an electronic automatic pipettor and further manually pipetting, if necessary, followed by seeding into fresh Matrigel domes in 24-well tissue culture plates (Corning 3526). Propagation was performed at 1:3-6 splitting ratio once every 5-7 days. For the first 4 days after re-seeding, the complete growth media was supplemented with an additional 5 µM Y-27632 (Sigma, Y0503).

### Derivation of STB^out^ TOs by suspension culturing

To generate STB^out^ TOs, mature STB^in^ organoids cultured as described above were first released from Matrigel domes using cell recovery solution (Corning, 354253) on ice with constant rotating at high speed (>120 rpm) for 30∼60 min, pelleted, washed one time with basal media (Advanced DMEM/F12 + 1% P/S + 1% L-glutamine + 1% HEPES) and resuspended in complete growth media supplemented with 5 µM Y-27632. Organoids were then carefully transferred using FBS pre-coated wide orifice p200 pipette tips (Fisher Scientific, 02-707-134) into an ultra-low attachment 24-well plate (Corning, 3473). One dome containing ∼ 500 organoids units can be dispensed into up to 5 wells of a 24-well plate with < 100 organoids units per well. TOs were evenly distributed in the wells prior to culturing in a 5% CO2 37℃ incubator for suspension culture of 1-2 d. Constant orbital rotating was introduced into suspension culture to improve polarity reversal efficiency (Thermo Fisher, 88881103). Media was renewed daily, and any aggregates dissociated using a FBS pre-coated wide orifice p200 pipette tip.

### Single organoid STB^out^ suspension cultures in 96-well plates

To perform single organoid unit suspension culture, each individual mature STB^in^ TO unit was picked out from domes with a sterilized needle (BD, 305125), including removal of any trace Matrigel matrix without compromising organoid integrity using a light microscope. Isolated organoids were then placed into a U-bottom well of an Ultra-low attachment 96-well spheroid microplate (Corning, 4515) for 2d suspension culturing as described above.

### Collection of conditioned media

Conditioned media (CM) was collected from original STB^in^ in domes as described ^1^. To harvest CM from STB^out^ TOs in suspension culture, the suspension culture 24-well plate was tilted for ∼2 min to sediment organoids to one side of the well, then carefully aspirate the supernatant media without disturbing the bottom organoids. Parallel STB^in^ and STB^out^ TOs wells used for CM collection contained approximately same initial number of organoids for following analysis.

### STB^in^ and STB^out^ TOs total protein extraction and quantification

STB^in^ TOs in Matrigel domes were released and collected as previously described^1^, then total protein was extracted using RIPA buffer containing proteinase inhibitor and sonication at 10 amplitudes, after 10 min incubation on ice, centrifuge at high speed > 12000 g for 10 min, finally collect the supernatant as organoids lysate. For the STB^out^ TOs total protein extraction, same protocol was used except skipping the organoids releasing step. Total protein quantifications were performed using a BCA Protein assay kit (Pierce, 23227) according to the manufacturer’s instructions.

### Immunofluorescence microscopy

STB^in^ TOs were immunostained as described^1^. For staining of STB^out^ TOs in suspension, the same protocol described was used, but the releasing of organoids from Matrigel was omitted. The following antibodies or reagents were used: SDC-1 (Abcam, ab128936), hCG beta (Abcam, ab243581), ITGA6 (Invitrogen, MA5-16884), HLA-G (Abcam, ab52454 and ab283260), cytokeratin-19 (Abcam, ab9221), Alexa Fluor 488 Goat anti-Mouse IgG secondary antibody (Invitrogen, R37120). Alexa Fluor 488 Goat anti-Rabbit IgG secondary antibody (Invitrogen, R37116), Alexa Fluor 488 Goat anti-Rat IgG secondary antibody (Invitrogen, A11006), Alexa Fluor 594 Goat anti-Mouse IgG secondary antibody (Invitrogen, R37121), Alexa Fluor 594 Goat anti-Rabbit IgG secondary antibody (Invitrogen, R37117), Alexa Fluor 633 Goat anti-Mouse IgG secondary antibody (Invitrogen, A21052), Alexa Fluor 647 Goat anti-Rat IgG secondary antibody (Invitrogen, A21247), Images were captured using a Olympus Fluoview FV3000 inverted confocal microscope or a Zeiss 880 Airyscan Fast Inverted confocal microscope and contrast-adjusted in Photoshop or Fiji. Image analysis and generation of three-dimensional movies was performed using Imaris (version 9.2.1, Oxford Instruments).

### Histochemistry and immunostaining of organoid frozen sections

Organoid preparation and embedding, cryosectioning, H&E staining, and immunostaining were performed as previously described^10^, using an H&E stain kit (Abcam, ab245880) according to the manufacturer’s instructions or standard procedure for immunostaining. Briefly, the collected STB^in^ and STB^out^ TOs were fixed (4% PFA/1×PBS) and permeabilized (0.5% Triton X-100/1×PBS), then submerged in 20% sucrose solution overnight, then finally embedded into the 7.5% gelatin/10% sucrose embedding solution and stored at -80 ℃. Cryosectionion of organoids frozen blocks was performed using a cryotome (Leica, CM1950) at 10 µm thickness. For immunostaining of cryosections, frozen sections were warmed to room temperature, then immunostaining performed with the antibodies as described above. Images were captured on a Keyence BZ-X810 all-in-one fluorescence microscope and contrast-adjusted in Photoshop.

### Luminex assays

Luminex assays were performed using the following kits according to the manufacturer’s instructions: hCG Human ProcartaPlex Simplex Kit (Invitrogen, EPX010-12388-901), Bio-Plex Pro Human Inflammation Panel 1 IL-28A / IFN-λ2 (Bio-rad, 171BL022M), Bio-Plex Pro Human Inflammation Panel 1, 37-Plex (Bio-rad, 171AL001M), and Bio-Plex Pro Human Chemokine Panel, 40-Plex (Bio-rad, 171AK99MR2). Plates were washed using the Bio-Plex wash station (Bio-rad, 30034376) and read on a Bio-Plex 200 system (Bio-rad, 171000205). All samples from both polarity conditions (STB^in^ and STB^out^) were tested in duplicate, and each condition was performed with at least three biological replicates from unique placental tissue. All measurements were normalized to total protein of either STB^in^ or STB^out^ TOs wells, quantified as described above.

### Cytotoxicity Assay

Cytotoxicity assays were performed using the CytoTox96^®^ Non-Radioactive Cytotoxicity Assay kit (Promega, G1780) according to the manufacturer’s instructions. All samples were tested in duplicate, and each condition (STB^in^ or STB^out^) was performed with at least three biological replicates.

### Coat-seeding of STB^in^ and STB^out^ TOs onto round coverslips for patch clamp

To seed collected original STB^in^ TOs onto the round glass coverslips (VWR, 76305-514) pre-coated with thin layer of Matrigel (Corning, 356231), each round coverslip was evenly distributed with ∼ 40 μl of Matrigel and carefully transferred into each well of regular 24-well plate to polymerize in a 37 ℃ incubator for ∼ 20 min. Then, organoids were harvested as described above and evenly dispensed onto the Matrigel pre-coated surface of coverslips to settle down in a 5% CO2 37 ℃incubator for 3∼4 h to ensure that the majority of organoids attach onto the matrix coating of the coverslip. For the STB^out^ TOs coat-seeding, the same protocol described above was used except omitting the release of organoids from Matrigel domes.

### Patch clamp estimation of cell surface area

All results were recorded in whole-cell configurations using an Axopatch 200B amplifier (Molecular Devices) and the pClamp 10 software package (Molecular Devices). The glass pipettes were pulled from borosilicate capillaries (Sutter Instruments) and fire-polished using a microforge (Narishge) to reach a resistance of 2–3 MΩ. The pipette solution (internal) contained (in mM): 140 CsCl, 1 MgCl_2_, 10 HEPES, 0.2 EGTA. pH was adjusted to 7.2 by CsOH. The bath solution contained (in mM): 140 CsCl, 10 HEPES, 1 MgCl_2_. pH was adjusted to 7.4 by CsOH. All experiments were at room temperature (22–25°C). All the chemicals for solution preparation were obtained from Sigma-Aldrich. Once the whole cell configuration was established, a 10-mV voltage command was delivered to the cell from a holding potential of 0 mV. The corresponding capacitive current was recorded. Membrane capacitance of the cell was calculated using Clampfit software (Molecular Devices) based on the following equation, 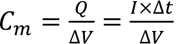, where Cm is the membrane capacitance, Q is the stored charge across the cell membrane, V is membrane voltage, I is current, and t is time. For the histogram plot, the bins (x-axis) were set as (pF): 0-20, 20-100, 100-200, 200-500, 500-1000, 1000-2000, 2000-5000 and 5000-10000. The bars on the histogram were set in the middle of each bin.

### Statistics and reproducibility

All experiments reported in this study have been reproduced using a minimum of three independent organoids lines derived from unique placental tissues. All statistical analyses were performed using Clampfit (Molecular Devices), Excel, or Prism software (GraphPad). Data are presented as mean ± SD, unless otherwise stated. Statistical significance was determined as described in the figure legends. Parametric tests were applied when data were distributed normally based on D’Agostino-Pearson analyses; otherwise, nonparametric tests were applied. For all statistical tests, p value <0.05 was considered statistically significant, with specific p values noted in the figure legends.

## Supporting information

**Supplemental Movie 1**: Three-dimensional image reconstruction of an STB^in^ trophoblast organoid (shown in Figure 2B, top) immunostained for SDC-1 (in green) and cytokeratin-19 (in red). DAPI-stained nuclei are shown in blue.

**Supplemental Movie 2**: Three-dimensional image reconstruction of an STB^out^ trophoblast organoid (shown in Figure 2B, bottom) immunostained for SDC-1 (in green) and cytokeratin-19 (in red). DAPI-stained nuclei are shown in blue.

**Supplemental Movie 3**: Three-dimensional image reconstruction of an STB^in^ trophoblast organoid (shown in Figure 2B, top) immunostained for ITGA6 (in green) and CGBs (in red). DAPI-stained nuclei are shown in grey.

**Supplemental Movie 4**: Three-dimensional image reconstruction of an STB^out^ trophoblast organoid (shown in Figure 2B, top) immunostained for ITGA6 (in green) and CGBs (in red). DAPI-stained nuclei are shown in grey.

## ACKNOWLEDGMENTS

This work was supported by the NIH grants R01AI145828 (C.B.C.) and DP2GM126898 (H.Y.). We also gratefully acknowledge the Duke Light Microscopy Core Facility for their technical support and assistance for this work.

## AUTHOR CONTRIBUTIONS

L.Y. and C.C. conceived the study, developed the methodology, and analyzed the data; P.L. and H.Y. performed patch clamping measurement, and analyzed the data; All authors participated in manuscript writing, review, and editing.

---

**Figure S1.**
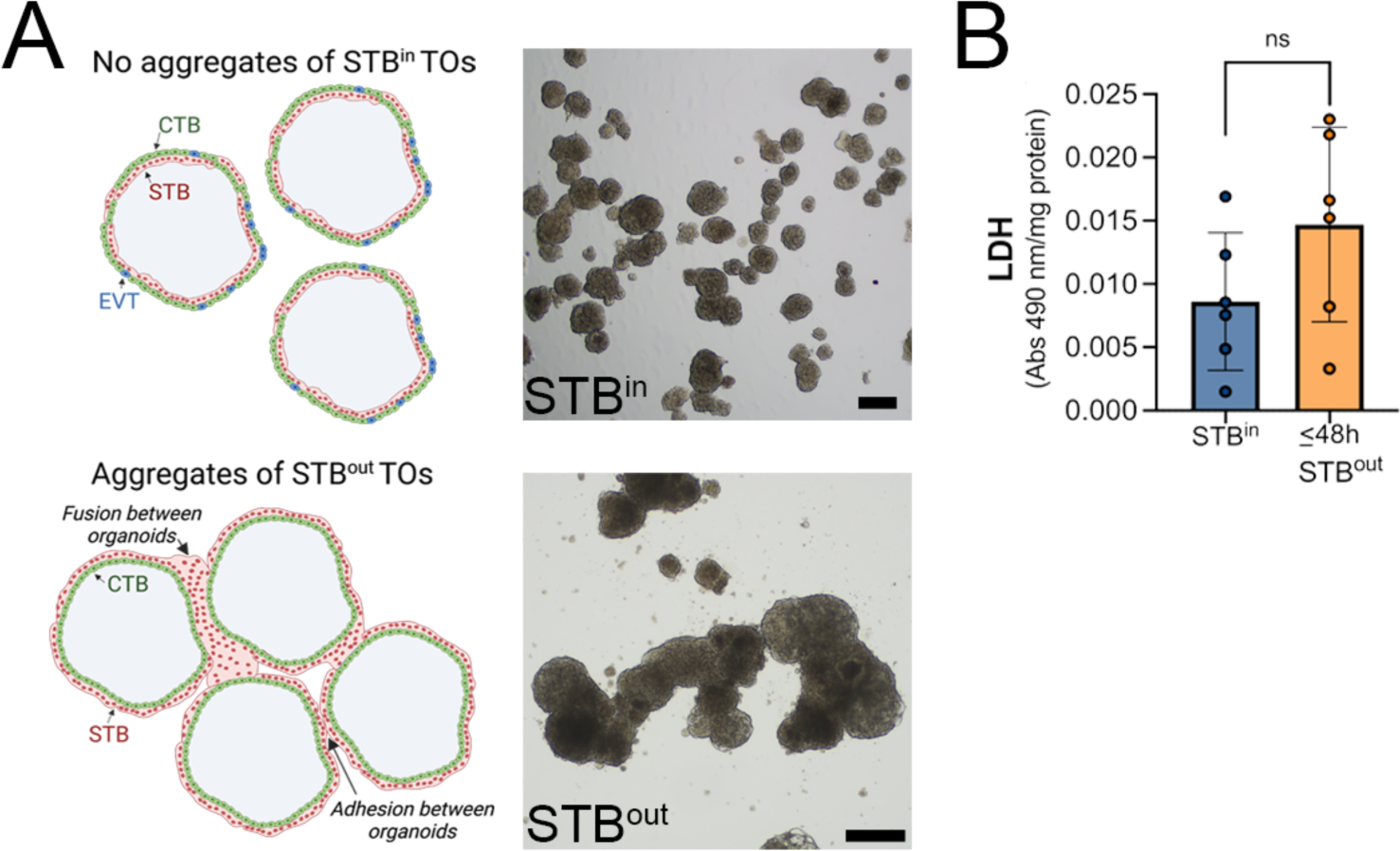
Evaluation of STB^in^ and STB^out^ TOs. **(A)**, Left, schematic of STB^in^ (top) or STB^out^ (bottom) TOs demonstrating the aggregation that can occurs in STB^out^ TOs that results from fusion of the STB and/or adhesion between organoid units. At right, brightfield images of STB^in^ (top) or STB^out^ (bottom) TOs demonstrating the extent of aggregation that can occur. Scale, 150μm (top) and 125μm (bottom). All schematics created using Biorender**. (B),** Levels of lactate dehydrogenase (LDH) in conditioned medium from STB^out^ TOs cultured for ∼48hrs. Data are shown as 490nm absorbance normalized to total protein. Data are shown as mean ± standard deviation with significance determined by a student’s t-test (ns, not significant). Symbols represent unique fields of organoids from individual replicates.

**Figure S2.**
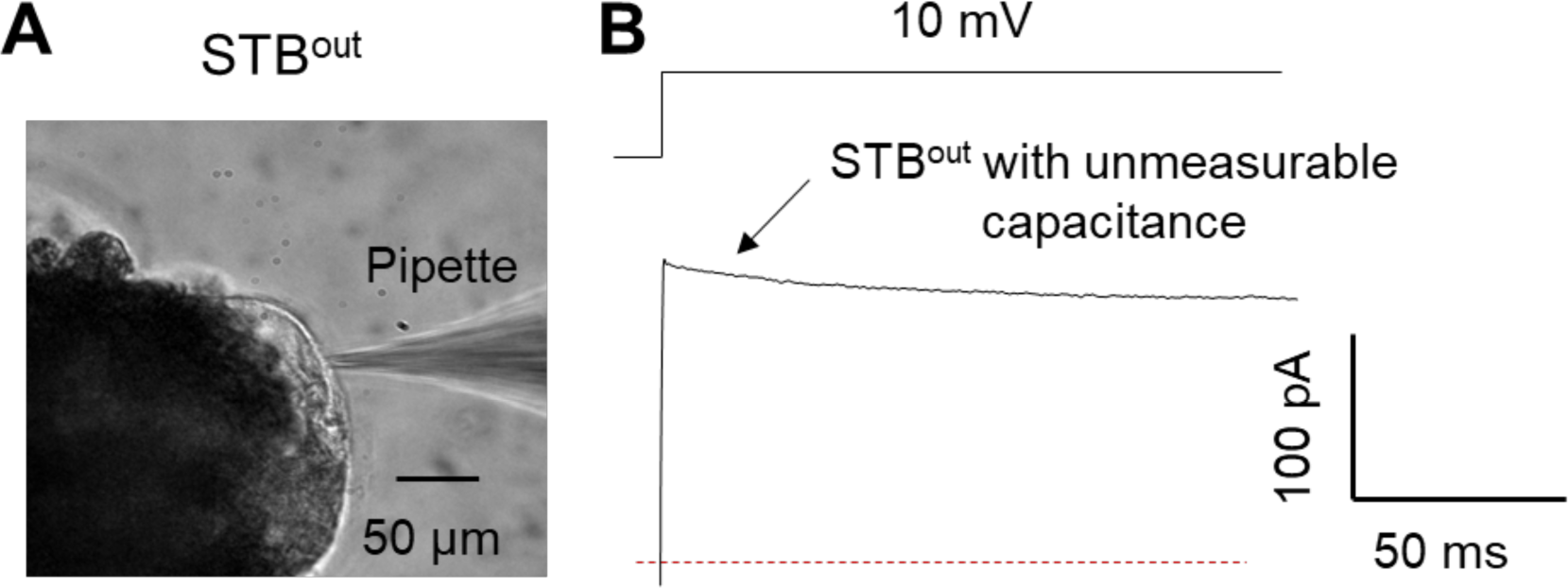
Patch clamp measurement of large syncytia from the surface of STB^out^ TOs. **(A)**, Representative brightfield image of a patch-clamped, extremely large syncytia from trophoblast organoids (TOs) growing under STB^out^ conditions. **(B),** Representative membrane test trace from large STB. Cell capacitance cannot be accurately measured due to the space clamp issue of large syncytia. Current was elicited by a test voltage pulse of 10 mV from a holding potential of 0 mV (top).

